# SSAlign: Ultrafast and Sensitive Protein Structure Search at Scale

**DOI:** 10.1101/2025.07.03.662911

**Authors:** Lei Wang, Xuchao Zhang, Yan Wang, Zhidong Xue

## Abstract

The advent of highly accurate structure prediction techniques such as AlphaFold3 is driving an unprecedented expansion of protein structure databases. This rapid growth creates an urgent demand for novel search tools, as even the current fastest available methods like Foldseek face significant limitations in sensitivity and scalability when confronted with these massive repositories. To meet this challenge, we have developed SSAlign, a protein structure retrieval tool that leverages protein language models to jointly encode sequence and structural information and adopts a two-stage alignment strategy. On large-scale datasets such as AFDB50, SSAlign achieves a two-orders-of-magnitude speedup over Foldseek in search, substantially improving scalability for high-throughput structural analysis. Compared to Foldseek, SSAlign retrieves substantially more high-quality matches on Swiss-Prot and achieves marked performance improvements on SCOPe40, with relative AUC increases of +20.2% at the family level and +33.3% at the superfamily level, demonstrating significantly enhanced sensitivity and recall. In sum, SSAlign achieves TM-align–comparable accuracy with Foldseek-surpassing speed and coverage, offering an efficient, sensitive, and scalable solution for large-scale structural biology and structure-based drug discovery.

## Introduction

Efficient identification of homologous proteins is fundamental to understanding biological function, reconstructing evolutionary relationships, and enabling structure-based drug discovery^1–5^. While sequence-based methods are widely used, they often struggle to detect distant homology. In contrast, the higher conservation of three-dimensional structures offers a powerful alternative for revealing functional and evolutionary connections. Recent breakthroughs in protein structure prediction have transformed the field. AlphaFold2 has led to the release of over 214 million predicted structures^6^, and ESMFold has contributed more than 617 million metagenomic protein models through the ESM Atlas^7^. With the advent of AlphaFold3^8^, both the scale and complexity of structural data continue to grow rapidly. This explosion of structural information presents unprecedented opportunities for biological discovery, while simultaneously raising a pressing challenge: how can we perform fast and scalable comparisons across hundreds of millions of protein structures without compromising accuracy? Addressing this question is key to unlocking the full potential of structural genomics.

Sequence-based approaches such as MMseqs2^9^, BLASTp^10^, and DIAMOND^11^ have long been the mainstay for large-scale homology detection, owing to their speed and scalability. However, relying solely on sequence similarity often fails to uncover remote evolutionary relationships, as protein tertiary structures are generally more conserved than primary amino acid sequences^12^. Structure alignment tools such as TM-align^13^ provide higher sensitivity for detecting distant homologs, but their computational cost is prohibitive. For example, searching a database of 100 million protein structures with a single query using TM-align can take over a month^13^. To balance sensitivity and efficiency, Foldseek encodes protein backbone geometry into a 20-letter structural alphabet, enabling rapid k-mer-based search and alignment^14^. While Foldseek is currently the fastest structural alignment method available, the accelerating growth of predicted protein structure databases—expected to reach tens of billions—poses significant computational challenges. At such scale, Foldseek remains time-consuming for large-scale structural searches. Moreover, its reliance on discretized structural representations may fail to capture intricate spatial interactions and subtle folding patterns, potentially leading to the omission of highly similar structures during the prefiltering stage.

Meanwhile, a novel research paradigm has emerged: Protein Language Models (PLMs). These models leverage advances in Transformer architectures to treat amino acid sequences as a form of “biological language,” learning contextual embeddings from large-scale natural protein sequence datasets. Such embeddings implicitly capture evolutionary information and structural constraints within sequences, providing a new avenue for remote homolog identification without reliance on sequence alignment—an area where traditional sequence comparison methods often fall short. This approach has demonstrated considerable potential in sequence retrieval tasks. For instance, Dense Homolog Retriever (DHR)^15^ employs embeddings generated by protein language models for dense vector similarity search, enabling efficient identification of remote homologs without alignment and outperforming traditional methods. However, DHR relies on large-scale supervised contrastive learning, and its performance on previously unseen, novel sequences remains limited. Nevertheless, DHR’s success validates the practical utility of the biological semantics encoded in PLM embeddings and provides an effective tool to overcome the limitations of sequence similarity. With the ongoing evolution of Transformer architectures, protein language models have continued to advance. Early models such as UniRep^16^ and ProtTrans^17^ have been widely applied in sequence modeling tasks, while the ESM family^18–20^ has progressively pushed the boundaries of representational capacity. Recent efforts have further integrated explicit structural information into PLM embeddings—for example, SaProt^21^ generates structure-aware embeddings by discretizing three-dimensional features, while ESM-3^20^ integrates sequence, structure, and function into unified multimodal representations. Given its efficiency and representational strength, SaProt serves as the backbone of our proposed system for structural retrieval.

We present SSAlign, a high-throughput structural retrieval system that integrates the SaProt model with dense vector search to identify structural homologs at scale. SSAlign encodes protein structures into fixed-length embeddings optimized for structural separability in latent space. Candidate structures are retrieved using approximate nearest neighbor search, followed by a lightweight two-stage filtering procedure to refine precision. As illustrated in Fig. 1, the pipeline begins with parallel encoding of input protein structures using Foldseek’s structural encoder—which discretizes backbone geometry and residue context into 3Di tokens—and the SaProt protein language model, which generates deep sequence embeddings. These two representations are jointly refined by the Entropy Reduction Module (ERM), which performs spatial redistribution and dimensionality reduction to produce unified embeddings for both queries and database entries. A fast vector similarity search then identifies candidate homologs, which are further evaluated by the SS-score predictor to estimate alignment quality. For candidate pairs below a predefined confidence threshold, the SAligner module applies global alignment using an accelerated Needleman–Wunsch algorithm with a 3Di substitution matrix, yielding a final set of high-confidence structural matches. Fully parallelized across multi-GPU or CPU-only hardware, SSAlign completes queries against tens of millions of protein structures in seconds. Compared to Foldseek, SSAlign achieves a two-orders-of-magnitude speedup while substantially improving recall and sensitivity. On benchmarks such as SCOPe40, SSAlign increases family-level AUC by 20.2% and superfamily-level AUC by 33.3% relative to Foldseek, and retrieves more high-quality matches (delivers alignment quality—measured by TM-score and RMSD—comparable to TM-align) than Foldseek on the Swiss-Prot dataset. Notably, SSAlign is also effective in retrieving small peptides or simply folded proteins—cases where Foldseek and related tools often struggle—due to its fine-grained structural representation and more sensitive embedding space. By bridging deep representation learning with scalable, GPU-accelerated search infrastructure, SSAlign offers an efficient, accurate, and scalable solution for large-scale structural biology, function annotation, and structure-based drug discovery.

**Figure 1.**
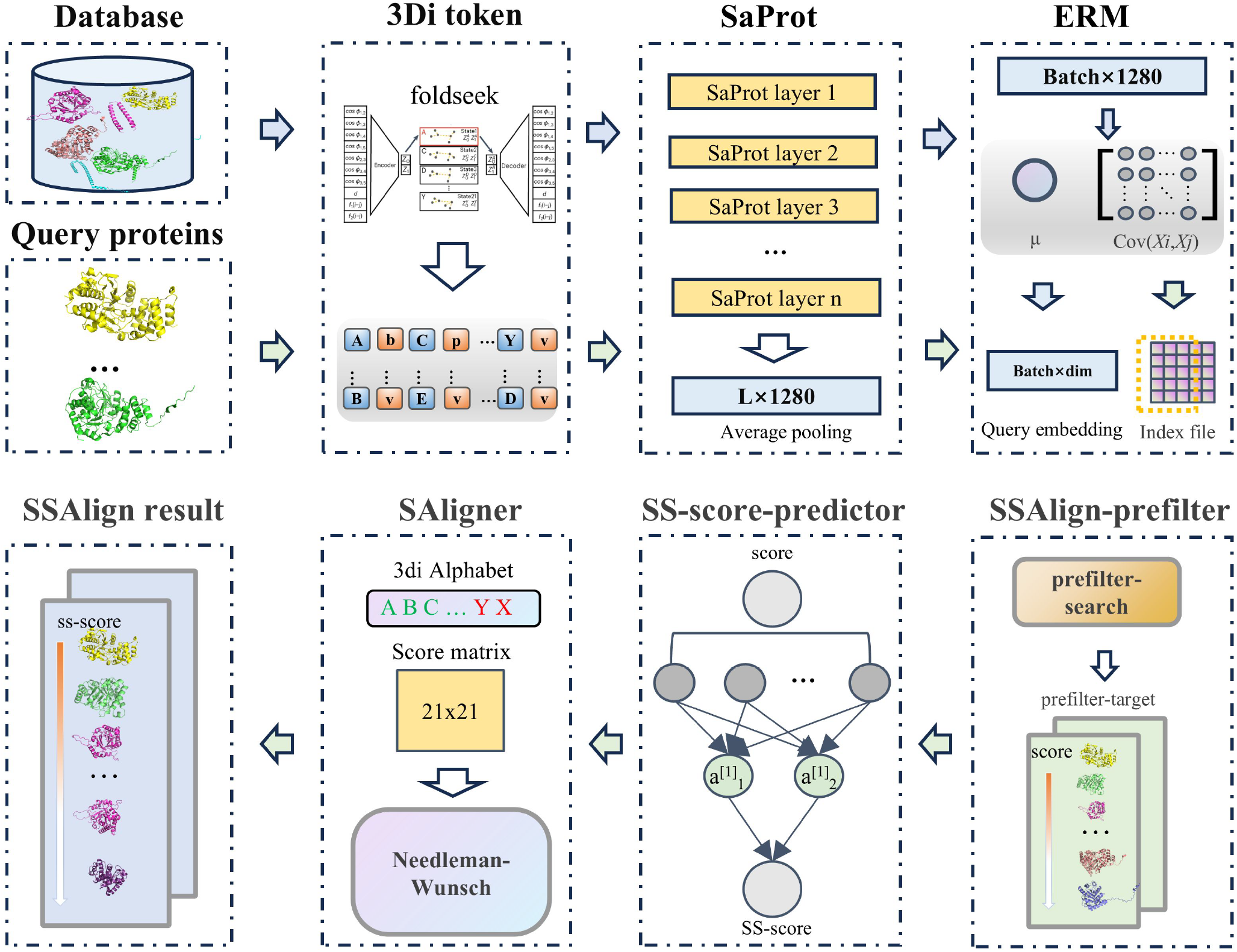
Overview of the SSAlign workflow. SSAlign employs a two-stage pipeline to efficiently search a set of query proteins against large structural databases. (a) Input protein structures are encoded in parallel using Foldseek’s structural encoder—which discretizes backbone geometry and residue context into 3Di tokens—and the Saprot protein language model—which generates deep sequence embeddings. These two representations are jointly refined by the Entropy Reduction Module (ERM), which performs spatial redistribution and dimensionality reduction to produce unified embeddings for both queries and database entries. A fast vector similarity search is then performed to identify candidate homologs. (b) The SS-score predictor maps similarity search metrics to estimated structural alignment scores. For candidate pairs falling below a predefined confidence threshold, the SAligner module applies a global alignment using an accelerated Needleman–Wunsch algorithm with a 3Di substitution matrix, yielding a final set of high-confidence structural matches.

## Results

To comprehensively evaluate the performance of SSAlign, we conducted a series of benchmark experiments designed to assess its computational efficiency, alignment accuracy, and robustness in detecting challenging protein relationships. Specifically, we evaluated: (1) its scalability and efficiency in large-scale structure search tasks; (2) its alignment accuracy and sensitivity compared to the state-of-the-art structure-based methods; and (3) its effectiveness in identifying homologous proteins with repetitive or highly simplified folds—cases that Foldseek often fails to capture. The following subsections detail the results of these evaluations.

### Order-of-magnitude acceleration in structure search

We evaluated the speed performance on the AFDB50 dataset by randomly selecting 1,000 proteins as query set, comparing SSAlign against Foldseek under controlled conditions.The benchmark setup utilized a high-performance computing system equipped with three NVIDIA RTX A6000 GPUs, each with 48GB of memory, two Intel Xeon Gold 6133 processors operating at 2.50GHz, and 256GB of system RAM. The specific parameters used were as follows:

- CPU execution time was measured by running SSAlign on CPU with the parameters: `--prefilter_mode cpu- -prefilter_target 2000 --prefilter_threshold 0.3 --max_target 1000 --nproc 64 -- mode 1`.
- GPU execution time was measured by running SSAlign on GPU with the parameters: `--prefilter_mode gpu- -prefilter_target 2000 --prefilter_threshold 0.3 --max_target 1000 --nproc 64 -- mode 1`.
- Foldseek was executed using its default parameters, with the addition of `--threads 64` to match the CPU core count used by SSAlign.
- TM-align, due to TM-align’s prohibitive computational cost (estimated execution time >1 month for this test scale), it was excluded from timing comparisons.

As demonstrated in Table 1, SSAlign achieves a remarkable two-orders-of-magnitude speedup compared to the baseline Foldseek on the AFDB50 dataset. While Foldseek requires 325,081 seconds (approximately 90 hours) to complete the 1,000 query tasks, SSAlign finishes the same workload in just 3,150 seconds on a standard CPU and 2,290 seconds with GPU acceleration. This corresponds to a 103*×* speedup on CPU and a 142*×* speedup on GPU, respectively. These results indicate that SSAlign effectively overcomes the computational bottlenecks of large-scale structural search, slashing the runtime from days to less than an hour. Notably, the massive acceleration on the CPU configuration confirms that this efficiency stems primarily from the optimized hierarchical algorithmic design rather than a strict reliance on high-end hardware acceleration.

**Table 1.** Runtime comparison and speedup of SSAlign versus Foldseek on the AFDB50 dataset (1,000 queries).

| Method / Stage | Time (Seconds) | | Speedup (×) | |
| --- | --- | --- | --- | --- |
|  | CPU | GPU | CPU | GPU |
| **Foldseek** (easy-search) | 325,081 | – | 1.0 | – |
| **SSAlign (Ours)** |  |  |  |  |
| Preload | 634 | 898 |  |  |
| Prefilter | 1,645 | 521 |  |  |
| SAligner | 871 | 871* |  |  |
| **Total** | **3,150** | **2,290** | **103.2** | **141.9** |

\*The SAligner stage is executed on the CPU in both configurations.

The stage-wise breakdown further elucidates the source of this efficiency and the trade-offs involved in GPU utilization. The GPU provides the most substantial gain in the `Prefilter` stage, reducing the runtime from 1,645 seconds to 521 seconds (∼ 3.2*×* faster), although this is partially offset by a slight increase in the `Preload` time (898 seconds vs. 634 seconds) due to the I/O overhead associated with transferring massive index data to GPU memory. Crucially, the fine-grained `SAligner` step, which is executed on the CPU in both configurations, consumes only 871 seconds, demonstrating that the pre-filtering mechanism acts as a highly effective sieve that narrows down the search space to a minimal fraction of the database. Consequently, SSAlign makes high-throughput structural analysis feasible on consumer-grade hardware, offering a highly accessible solution for metagenomic-scale studies.

### Comparative Benchmarking of SSAlign Against Foldseek and TM-align

We systematically benchmarked SSAlign, Foldseek, and TM-align on three widely used structural alignment datasets: Swiss-Prot, SCOPe40 and AFDB50. The term SSAlign-prefilter refers to the first-stage output of SSAlign prior to final scoring. Detailed evaluation procedures are described in the Datasets and Benchmark section.

#### Benchmarking on the Swiss-Prot Dataset

On the Swiss-Prot dataset, we randomly selected 100 query proteins and conducted a reference-free, unbiased evaluation by analyzing the cumulative TM-score and root-mean-square deviation (RMSD) across retrieved protein pairs. As shown in Fig.2a and 2c), when ranked by TM-align-derived avg_TM-score, SSAlign (including its prefilter module) achieved comparable cumulative avg_TM-score and RMSD performance to TM-align, while outperforming Foldseek, which showed a noticeable drop in cumulative TM-score—indicating its tendency to miss structurally similar targets. When results were sorted based on each tool’s internal scoring metric (E-value or SS-score, Fig.2b and 2d), TM-align remained the most effective, with Foldseek slightly outperforming SSAlign in ranking quality, albeit with limited impact on overall retrieval coverage. Importantly, SSAlign returned a significantly greater number of matched protein pairs than Foldseek. These additional results maintained high cumulative TM-scores and low RMSDs, indicating that SSAlign can improve recall without compromising alignment accuracy. A further comparison of method-specific match sets (i.e., pairs uniquely identified by one method; Fig.3a–d) demonstrated that SSAlign identified more high-quality alignments (avg_TM-score ≥ 0.5), with better structural coherence as reflected by higher cumulative avg_TM-scores and lower RMSDs. These findings highlight SSAlign’s enhanced sensitivity in detecting remote homologs and its ability to retrieve biologically relevant alignments beyond the scope of other methods.

**Figure 2.**
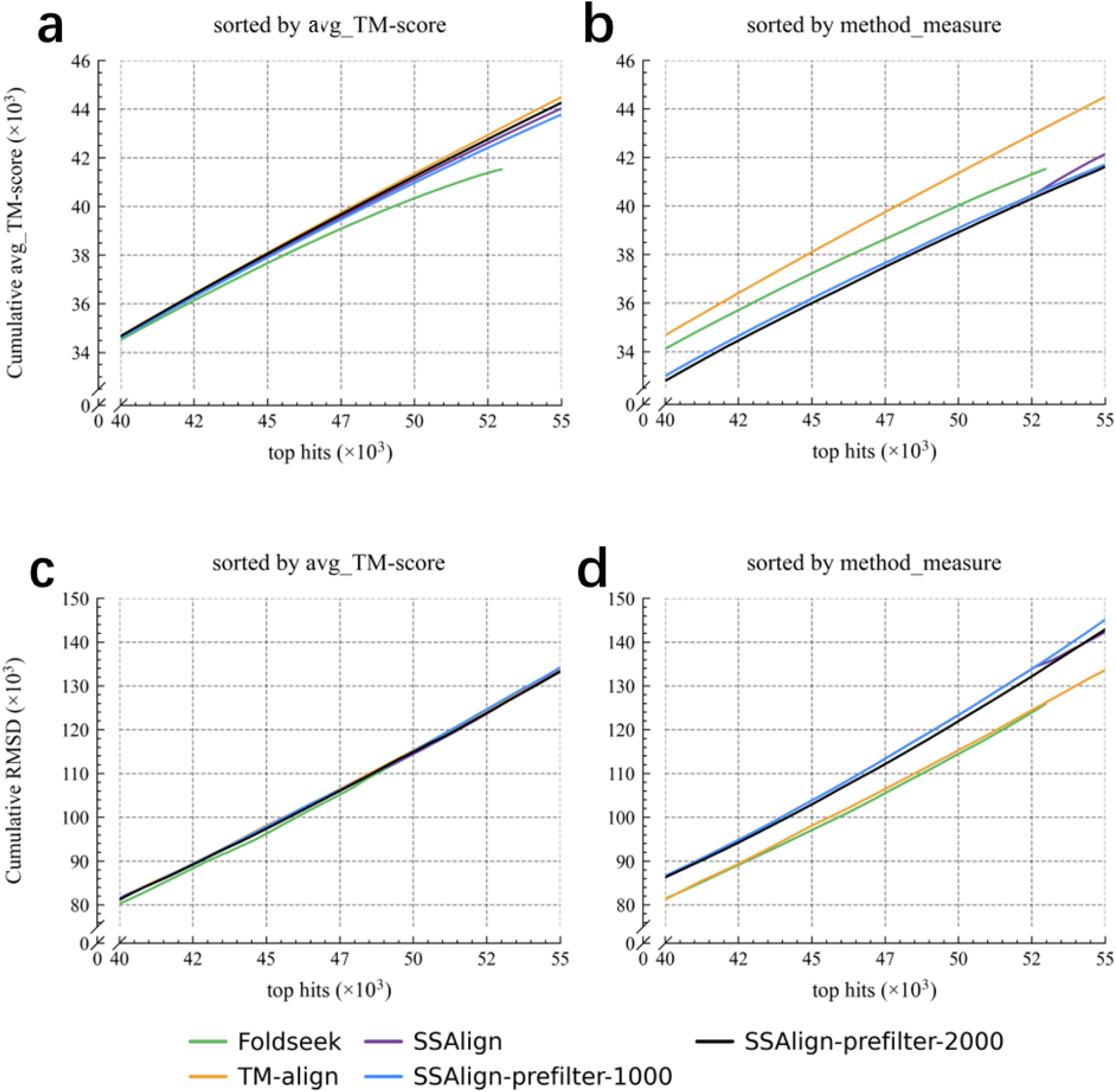
Comparative performance of structural alignment methods on the Swiss-Prot dataset. This benchmark utilizes the Swiss-Prot dataset comprising 100 query proteins against a database of 542,378 entries.(a-c) Display comparative results of five methods (ordered by TM-align-derived avg_TM-score), illustrating the cumulative avg_TM-score versus RMSD relationships for top-ranked protein pairs. (b-d) present results sorted by tool-specific metrics (E-value or SS-score). For each query protein, we performed exhaustive pairwise database searches using TM-align, applying an avg_TM-score ≥ 0.5 threshold for TM-align’s result. Consequently, TM-align consistently maintains optimal cumulative scores. The Foldseek curve truncation reflects its fewer returned results compared to other methods.

**Figure 3.**
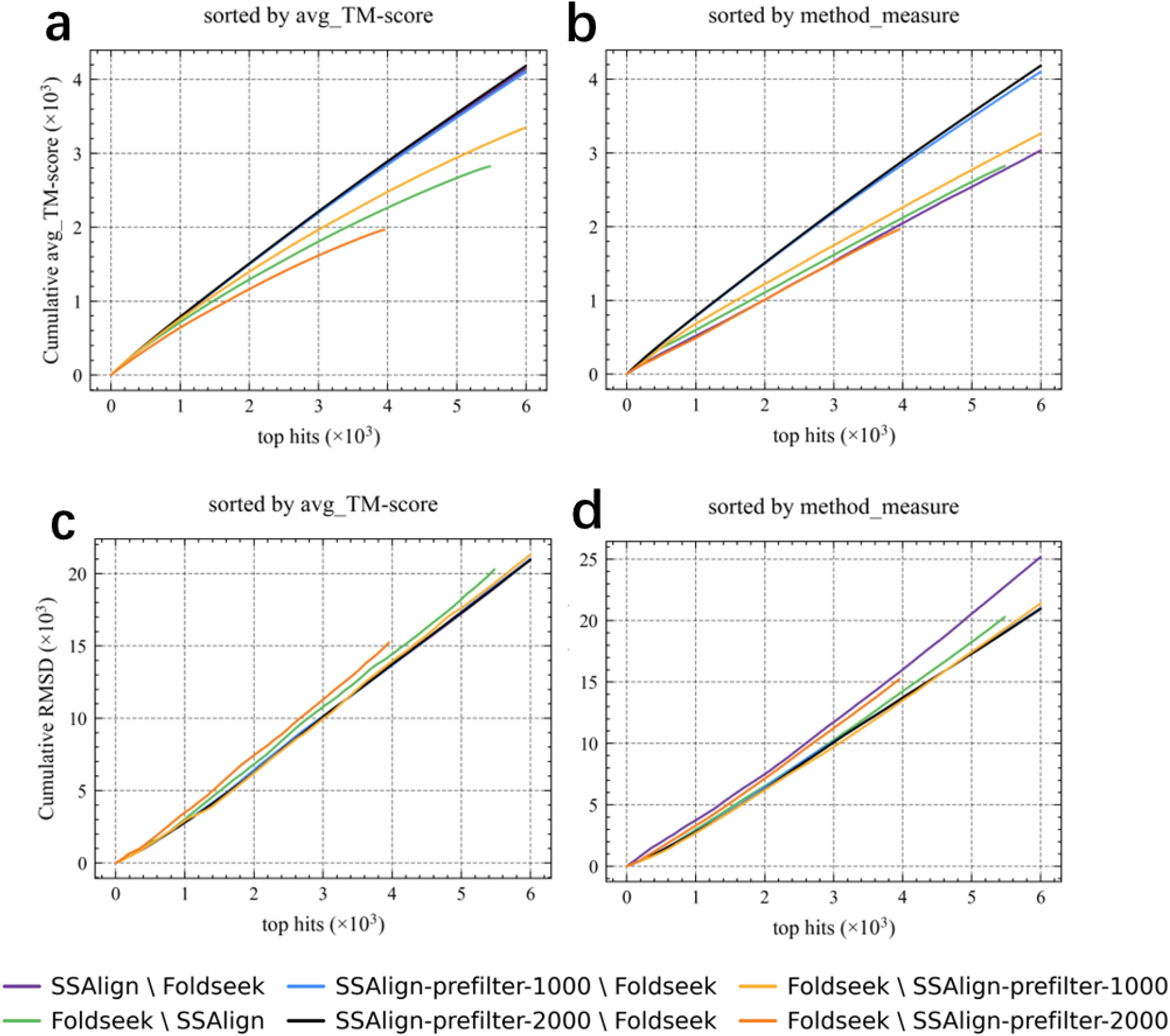
Structural comparison of differential matches between Foldseek and SSAlign on the Swiss-Prot dataset. This benchmark continues to employ the Swiss-Prot dataset comprising 100 query proteins. We examined the differential sets between hits identified by Foldseek and those by SSAlign/SSAlign-prefilter. (a-c) present comparative results of six differential sets (ordered by avg_TM-score), illustrating the cumulative avg_TM-score versus RMSD relationships for protein pairs within these differential sets; while (b-d) display results sorted by tool-specific metrics (E-value or SS-score). Notably, the cumulative avg_TM-score results conclusively demonstrate that SSAlign (including prefilter) can retrieve significantly more homologous protein pairs with high structural similarity.The legend annotates method-specific protein pairs (i.e., hits exclusively identified by one method).

#### Benchmarking on the SCOPe40 Dataset

We next assessed homology detection performance on the SCOPe40 dataset, which provides hierarchical family and superfamily annotations based on structural similarity—making it an ideal framework for evaluating structural alignment sensitivity. As shown in Fig.4a–b, SSAlign and its prefilter module outperformed Foldseek at both classification levels, identifying a larger number of true positive (TP) matches. The prefilter stage alone exhibited strong discriminative power, confirming the effectiveness of the embedding space and similarity filtering strategy. Although SSAlign returned fewer total hits than TM-align—owing to TM-align’s exhaustive scoring and tendency to yield redundant high-scoring matches—SSAlign achieved comparable precision–recall performance (Fig.4c–d) and surpassed Foldseek in recall. Quantitatively, SSAlign improved the area under the curve (AUC) at the family level from 0.213 (Foldseek) to 0.256 (+20.19%), and at the superfamily level from 0.186 to 0.248 (+33.33%), demonstrating robust and consistent gains in retrieval performance.

**Figure 4.**
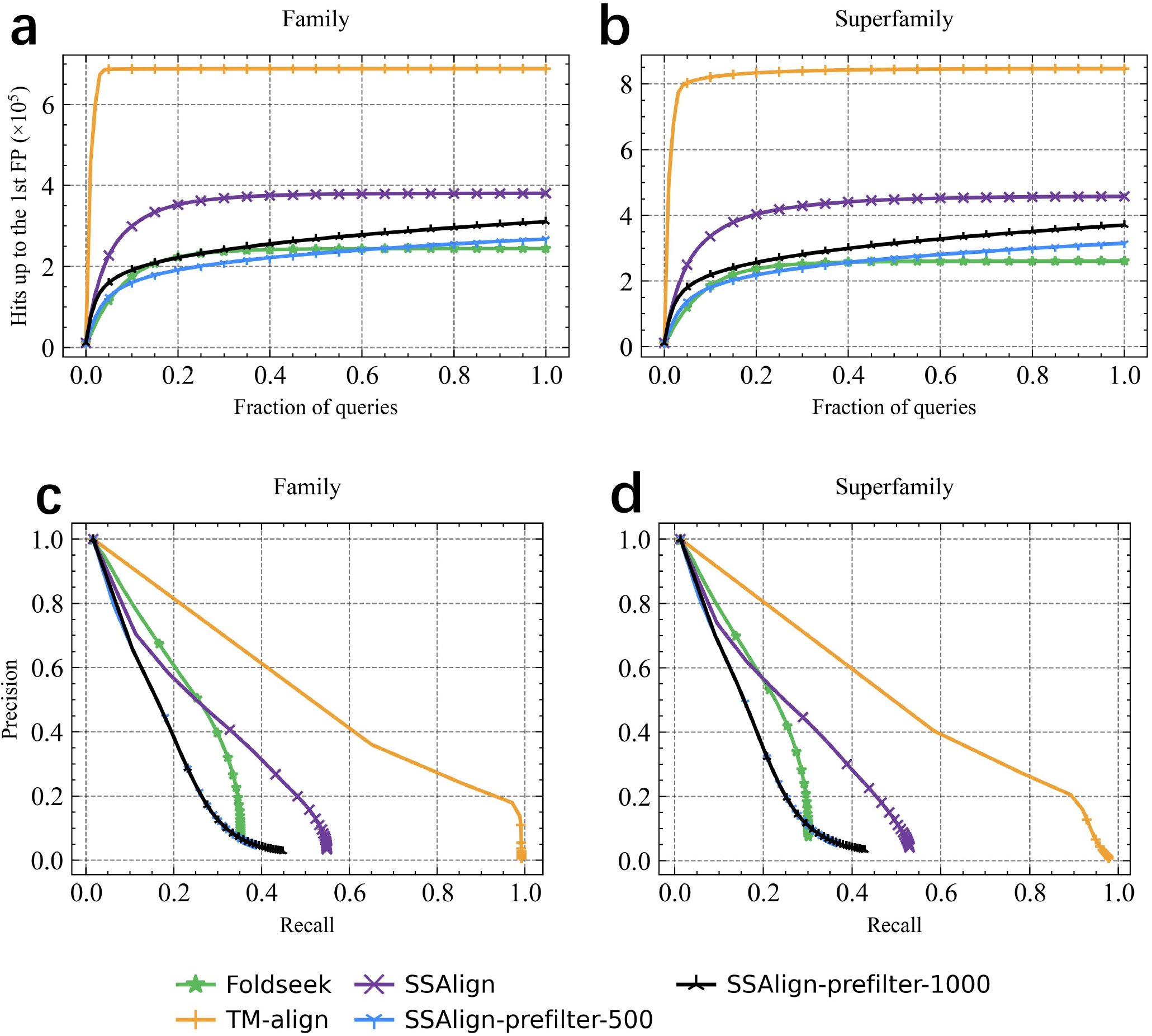
All-against-all performance evaluation on the SCOPe40 dataset. For family and superfamily level recognition, we defined true positives (TP) as : proteins belonging to the same family/superfamily, or those with avg_TM-score ≥ 0.5,False positives (FP) were classified as matches from different folds with avg_TM-score *<* 0.5.(a-b) Hits was calculated as the proportion of true positives (TPs) in the ranked list until the first false positive (FP) occurrence across different methods. (c-d) Precision-recall curves demonstrate the tools’ accuracy and recall performance. For TM-align, we performed all-to-all searches and sorted results by avg_TM-score. With each query protein returning 11,211 results, this approach achieved perfect recall.

#### Benchmarking on the AFDB50 Dataset

For benchmarking the runtime performance, we executed 1,000 query tasks on the AFDB50 dataset using SSAlign and Foldseek, respectively. Given the computational overhead of calculating the TM-score and RMSD metrics, we selected the 100 queries for which Foldseek returned the largest number of hits. A reference-free, unbiased evaluation was then performed by analyzing the cumulative avg_TM-score and RMSD across the retrieved protein pairs. As shown in Fig. 5a and 5d, when sorted by the avg_TM-score calculated from TM-align and with consistent top hits, SSAlign (including SSAlign-prefilter-2000) outperformed Foldseek in both cumulative avg_TM-score and RMSD performance. When the results were ranked according to each tool’s own scoring metric (E-value or SS-score, Fig. 5b and 5e), SSAlign exhibited the best performance, benefiting from the re-ranking effect of SAligner. Fig. 5c and 5f further show that the results returned by SSAlign achieved an average improvement of approximately 10% in avg_TM-score, alongside an average reduction of about 4.6% in RMSD. In the analysis of method-specific match sets, Fig. 6a–f reveals that although the set unique to Foldseek (Foldseek SSAlign) is larger than that unique to SSAlign (SSAlign Foldseek), the uniquely identified matches by SSAlign demonstrate significantly superior scores under the same top hits condition. Regarding additional hits unique to each method, the novel hits retrieved by SSAlign still maintain higher structural consistency: compared to the hits unique to Foldseek, they show an approximately 16.9% increase in avg_TM-score and an approximately 6.3% decrease in RMSD.

**Figure 5.**
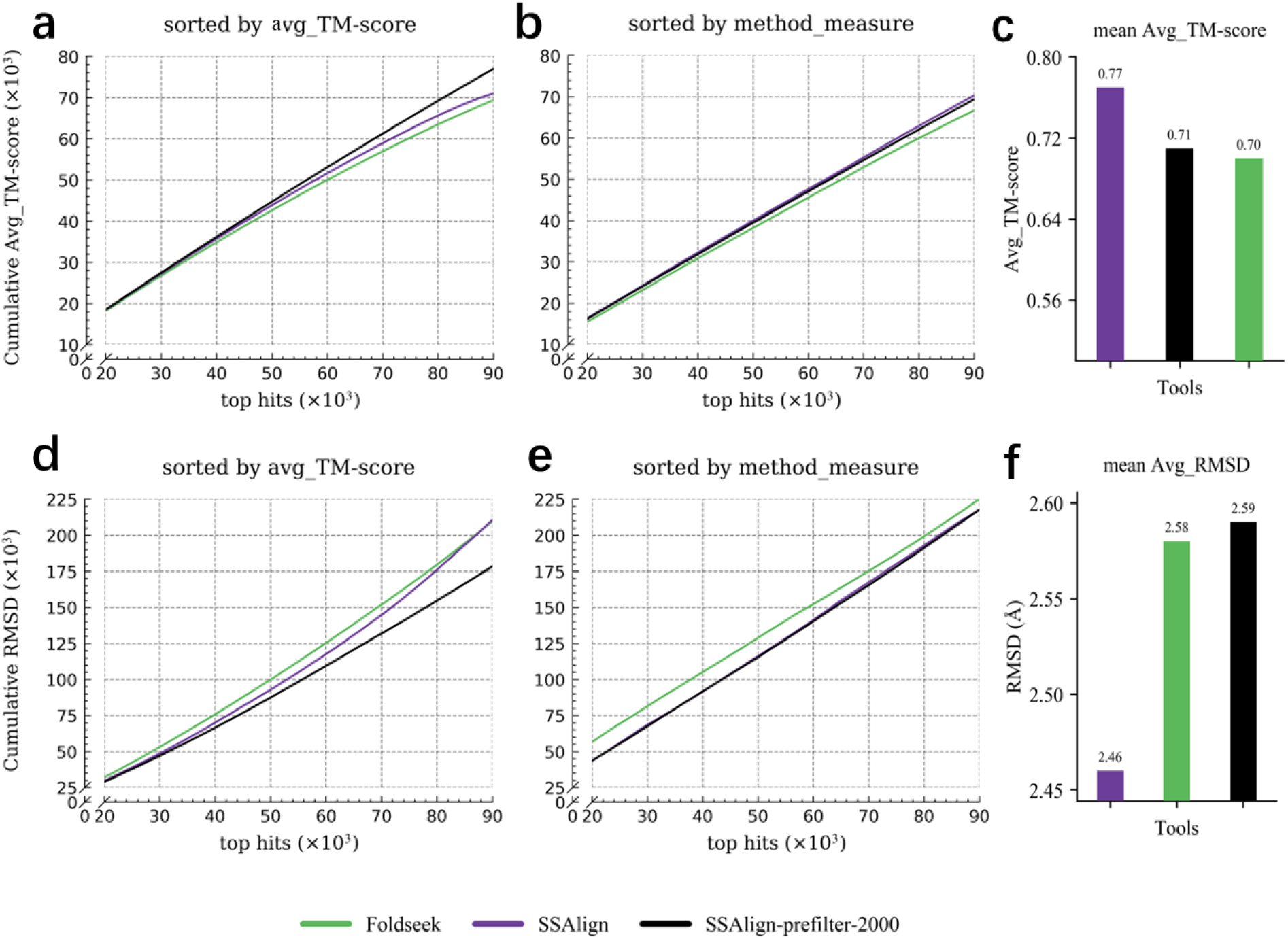
Comparative performance of structural alignment methods on the AFDB50 dataset. This benchmark utilizes the AFDB50 dataset comprising 100 query proteins against a database of 53,665,860 entries. (a–d) show the comparative results of three methods (sorted by TM-align-derived avg_TM-score), illustrating the cumulative avg_TM-score versus RMSD relationships for top-ranking protein pairs. (b–e) present the results ranked by tool-specific metrics (E-value or SS-score). Truncation of the curves reflects the relatively smaller number of results returned by the corresponding method. (c–f) display bar charts of the average scores for the results returned by each tool across the 100 tasks.

**Figure 6.**
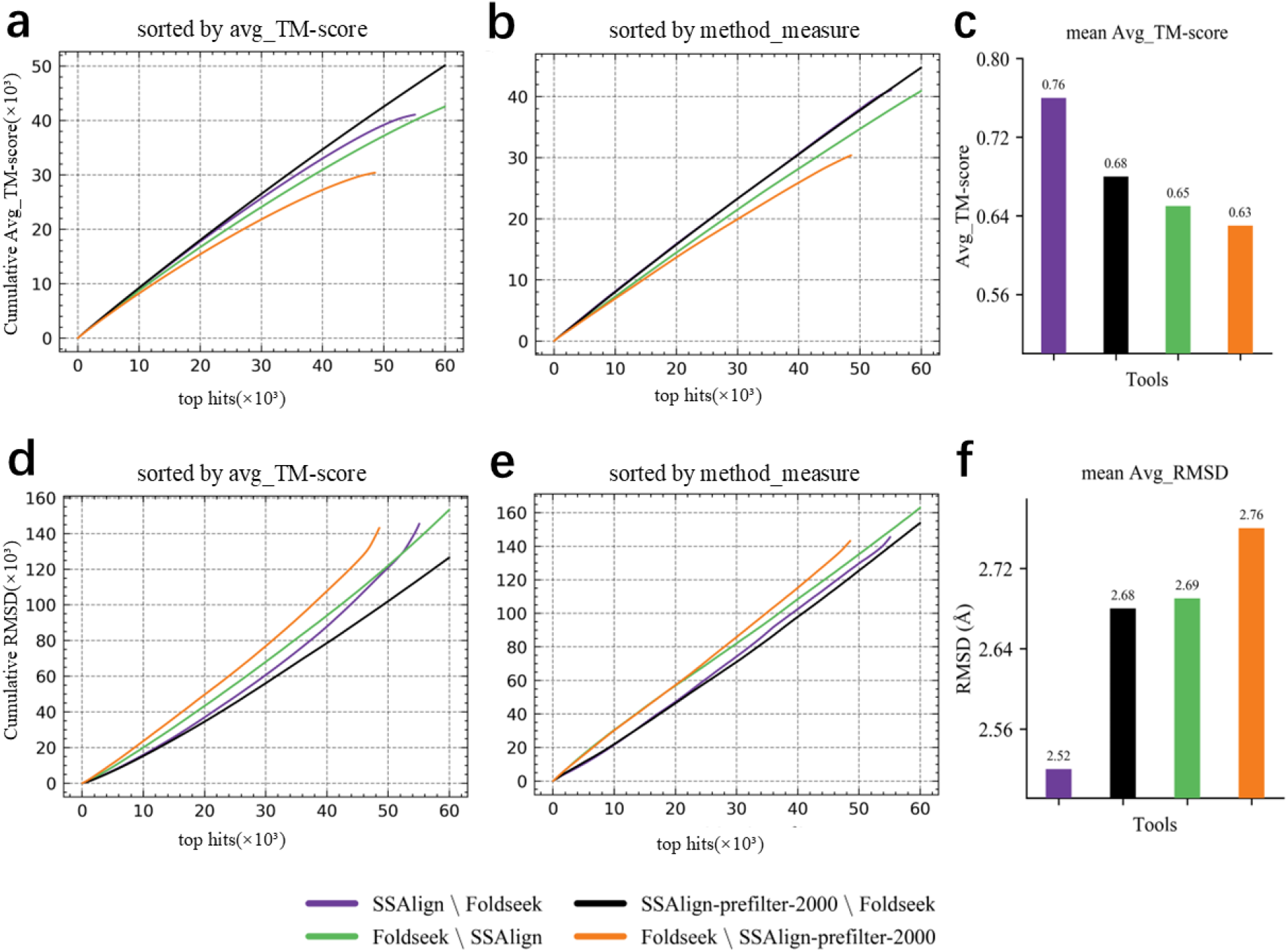
Structural comparison of differential matches between Foldseek and SSAlign on the AFDB50 dataset. This benchmark continues to employ the AFDB50 dataset containing 100 query proteins. We examined the differential sets between the hits identified by Foldseek and those identified by SSAlign/SSAlign-prefilter-2000. (a–d) present the comparative results of six differential sets (sorted by avg_TM-score), illustrating the cumulative avg_TM-score versus RMSD relationships for protein pairs within these differential sets; whereas (b–e) show the results sorted by tool-specific metrics (E-value or SS-score). (c–f) display bar charts of the average scores for the differential sets of results returned by each tool across the 100 tasks. The legend annotates method-specific protein pairs (i.e., hits exclusively identified by one method).

### SSAlign accurately detects simple fold proteins missed by Foldseek

Proteins composed of repetitive structural units, which we term “simple fold proteins” have long posed significant challenges in structural alignment. Foldseek achieves rapid structure search and comparison by compressing three-dimensional contact maps into one-dimensional 3Di sequences combined with an MMseqs2-based alignment approach. However, this encoding scheme inherently limits Foldseek’s sensitivity: the 3Di sequences use only 20 discrete states to represent contact environments and rely solely on nearest-neighbor contacts. As a result, simple fold proteins generate highly repetitive and monotonous 3Di patterns that are difficult to distinguish. This homogeneity often leads Foldseek’s k-mer prefiltering to discard true homologs containing repetitive motifs.

To systematically evaluate this limitation, we identified 13 pairs of simple fold proteins with clearly repetitive structural fragments within the SCOPe40 database. Despite their confirmed presence in the database, Foldseek failed to retrieve any matches for these proteins (Supplementary Table 1). By contrast, SSAlign, leveraging a more nuanced structural alignment strategy, successfully recovered the majority of these homologous relationships, demonstrating markedly improved sensitivity and specificity in recognizing simple fold architectures.

Furthermore, antimicrobial peptides (AMPs)—small, naturally occurring bioactive polypeptides—often adopt simple *α*-helical conformations characterized by repetitive and low-complexity structures with minimal sequence conservation^22^. These features align closely with our definition of simple fold proteins. We analyzed seven representative AMPs from the Swiss-Prot database (P84381–P84387) and found that Foldseek failed not only to detect homologs but also to retrieve self-alignments (Supplementary Table 2). In contrast, SSAlign successfully identified most homologous pairs. As shown in Fig. 7, using the antimicrobial peptide P84382 (XT-2) as the query, the top-ranked SSAlign hits display a highly consistent short *α*-helical conformation in PyMOL superposition. These hits also share a coherent biological context in UniProt annotations, being amphibian host-defense antimicrobial peptides secreted by skin glands and associated with the gastrin/cholecystokinin family / Magainin subfamily; P84382 is annotated as strongly active against *S. aureus* but weaker against *E. coli*. Together, this example indicates that SSAlign can reliably recover cross-species homologous candidates in low-complexity, repetitive-motif (simple-fold) regimes, where discrete 3Di k-mer prefiltering may miss true homologs. Detailed multi-parameter performance evaluations for these AMPs are provided in Supplementary Tables 3–9. Supplementary Table 10 lists additional AMPs’3Di sequences in Swiss-Prot that showed poor retrieval performance with Foldseek. In summary, SSAlign addresses key shortcomings of existing structure search methods in handling simple fold proteins and provides robust support for detecting functionally related but sequence-diverse structural units. This advancement holds significant implications for understanding the evolutionary relationships and functional diversity within complex protein families.

**Figure 7.**
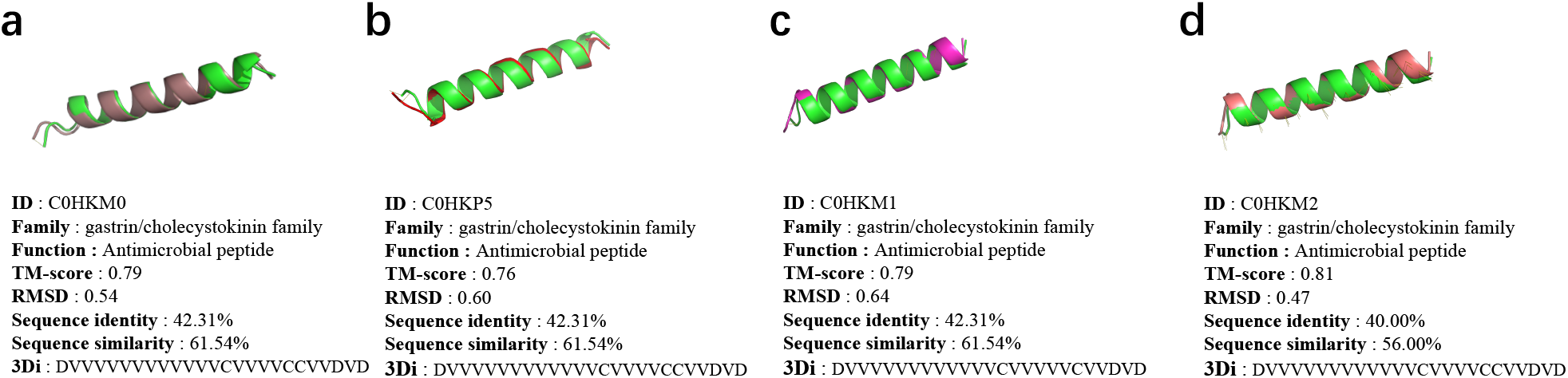
Structural superposition of the top-ranked SSAlign hits (C0HKM0, C0HKP5, C0HKM1, and C0HKM2) with the AMP query P84382 (XT-2). All hits share a conserved short *α*-helical fold and display high structural similarity (RMSD ≈ 0.47–0.65 Å; avg. TM-score ≈ 0.764–0.814).

## Discussion and Conclusions

SSAlign is a novel and highly efficient protein structure retrieval system that presents a significant advance in large-scale structural comparison. Its effectiveness arises from a synergistic architecture combining a powerful protein language model, a unique embedding optimization module, and rapid vector search technology. This system addresses key bottlenecks in computational speed and memory usage, offering a powerful alternative to existing methods like Foldseek for high-throughput structural bioinformatics.

A critical innovation of SSAlign is the Entropy Reduction Module (ERM), which significantly enhances retrieval recall by optimizing the initial protein embeddings. The ERM provides a computationally efficient solution to the problem of anisotropic embedding distribution, where certain vector dimensions can disproportionately influence similarity scores. By applying a simple linear transformation, the ERM decorrelates these dimensions and normalizes their variance, creating a more isotropic embedding space. This ensures that similarity calculations are more balanced and reliable, thereby boosting the discriminative power of the embeddings and directly leading to a higher recall rate for relevant structures during the critical initial screening phase.

Building upon this substantial improvement in retrieval accuracy, SSAlign also delivers exceptional computational speed. SSAlign demonstrates a substantial performance increase, completing large-scale queries in a dramatically shorter time—representing a two-orders-of-magnitude speedup compared to the 90 hours required by Foldseek for 1,000 protein structures under similar conditions. This efficiency, combined with reduced memory overhead, greatly enhances the feasibility of conducting proteome-wide structural searches.

Beyond its speed, SSAlign shows strong sensitivity, particularly in handling challenging proteins with repetitive motifs or simplified folds. In such cases, where other methods may struggle, SSAlign consistently retrieves high-quality structural homologs, validated by high cumulative TM-scores and low RMSD values. While SSAlign does not yet match the precision of TM-align’s top-ranked alignments, its primary strength lies in its capacity for high-recall, large-scale initial screening. In practical research workflows, it can serve as a highly effective tool for this preliminary step, making the study of protein relationships more comprehensive and efficient. The technical framework of SSAlign provides a solid foundation for future development. Future work will focus on enhancing the SS-score predictor and refining the global alignment algorithm within the SAligner module to further improve alignment precision. The exploration of methods such as adaptive embedding dimensionality, perhaps in concert with further refinements to the ERM, may yield a more optimal balance between computational efficiency and search accuracy. Furthermore, this core technology could be extended to new areas, such as the structural retrieval of protein-protein complexes, holding significant promise for applications in structural biology, drug discovery, and protein design.

## Methods

### Structure-aware Protein language model

SaProt is a large-scale, high-performance protein language model (PLM) trained on extensive protein sequence and structural data. To incorporate structural information, we utilized the 3Di structural discretization module from Foldseek to convert 3D protein structures into discrete structural tokens, which were then combined with residue-level tokens to construct a structure-aware (SA) vocabulary. This vocabulary serves as input to a conventional PLM architecture, enabling unsupervised training on large-scale SA-token sequences and resulting in a structure-aware PLM.

With 650 million parameters, SaProt is the largest PLM to date trained on the most comprehensive protein structure dataset. Training was conducted over 3 months using 64 NVIDIA A100 80GB GPUs, with total computational cost comparable to that of ESM-1b^18^. Built upon the ESM architecture, SaProt enhances structural representation via SA-token–based pretraining and demonstrates strong performance across a range of downstream tasks. To determine the most informative representation, we compared embeddings from the final (33rd) layer with those obtained by concatenating the first and final layers. The final-layer embeddings consistently yielded better performance across our evaluations (Supplementary Table 11) and were therefore used in all subsequent experiments.

#### ERM

To enhance the applicability of SaProt-generated embeddings for similarity search in the SSAlign-prefilter stage, a common approach involves transforming the initial protein language model into a dual-encoder framework through contrastive learning to separately encode query and target proteins^15^. However, the substantial computational costs associated with this method often limit its widespread adoption. Another prevalent strategy employs flow-based models to calibrate the distribution of sentence vectors derived from BERT, which has been extensively utilized in textual semantic similarity retrieval. Nevertheless, as a vector transformation model, the inherent design of flow-based models necessitates simple inverse transformations at each layer to ensure tractable Jacobian determinant calculations, resulting in limited nonlinear transformation capacity per layer. In practice, only half of the variables undergo transformation at each layer, necessitating considerable model depth to achieve sufficient fitting capability, which in turn demands extensive computational resources. We developed a resource-efficient Entropy Reduction Module (ERM) that achieves performance comparable to the aforementioned methods. Specifically, the high-dimensional embeddings produced by SaProt exhibit significant inter-dimensional correlations within the original space, causing certain dimensions to disproportionately dominate similarity calculations (Equation 1) and compromise result accuracy. Additionally, the dimensions demonstrate heterogeneous scaling (i.e., unequal variances), enabling dimensions with larger scales to exert disproportionate influence on similarity computations. Collectively, these factors yield an elliptical distribution of embeddings in the original space, characterized by dense data concentrations along certain axes and sparser distributions along others. Such anisotropic distributions distort similarity measurements by overemphasizing specific directional components.

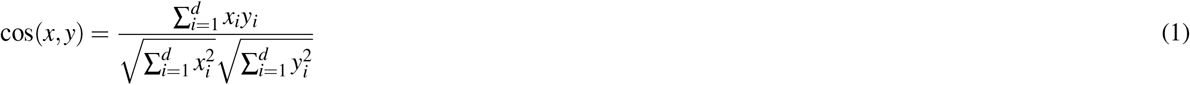

The ERM employs linear transformations to eliminate inter-dimensional correlations in the data (see Algorithm 1). This ensures that cosine similarity calculations more accurately reflect directional relationships between vectors by removing redundant information, while standardization balances the contribution of each dimension to similarity measurements. ERM converts the original elliptical embedding distribution into a spherical one, equalizing data density across all directions. This geometric transformation prevents bias in distance (or angular) calculations caused by anisotropic data distributions. Additionally, ERM alleviates the curse of dimensionality, where high-dimensional vectors tend to have similar pairwise distances, reducing the reliability of similarity comparisons. By optimizing the vector space geometry, ERM enhances the discriminability of vector distances, improving retrieval accuracy and efficiency.

##### Algorithm 1 ERM Workflow

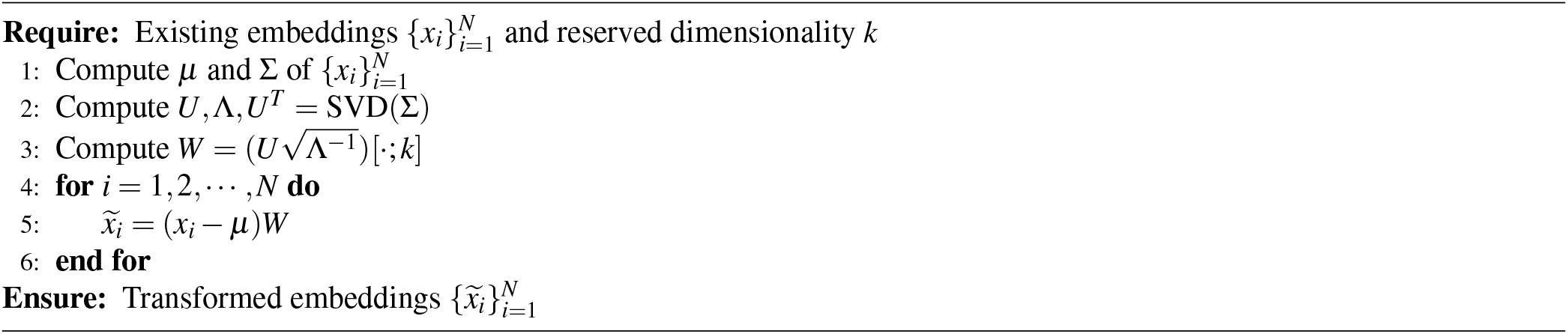

By retaining only the leading dimensions of the transformed embeddings, we achieve effective dimensionality reduction. Experimental results demonstrate that reducing embeddings to 512 dimensions not only maintains performance but even improves screening efficacy. To validate ERM’s effectiveness, we conducted tests on a randomly sampled subset from the Swiss-Prot dataset. As shown in Table 2, we compared the overlap rates between top-k results from SSAlign-prefilter and Foldseek using original embeddings and ERM-processed embeddings. At the full 1280 dimensions, ERM increases the overlap rate by approximately 32%, while the 512-dimensional embeddings achieve comparable performance.After applying ERM, reducing the embedding dimensionality from 1280 to 512 does not compromise retrieval effectiveness. Specifically, the overlap rates remain comparable at top-*k* = 1,000 and 2,000 (85.36/89.94 for 1280 dimensions vs. 83.70/89.36 for 512 dimensions), and at top-*k* = 4,000 the 512-dimensional embeddings even achieve a slightly higher overlap (92.86 vs. 92.41). Importantly, the reduced-dimensional embeddings substantially decrease memory usage and computational time (with an approximate 50% reduction in memory consumption) while preserving accuracy. Consequently, SSAlign adopts the ERM-processed 512-dimensional embeddings as the protein feature representation.

**Table 2.** Comparison of overlap rates between Foldseek and SSAlign-prefilter (with/without ERM module)

| Type | Dimension | Top-*k* | Overlap Rate (%) |
| --- | --- | --- | --- |
| without ERM | 1280 | 1,000 | 53.32 |
|  |  | 2,000 | 57.69 |
|  |  | 4,000 | 62.17 |
| with ERM | 1280 | 1,000 | 85.36 |
|  |  | 2,000 | 89.94 |
|  |  | 4,000 | 92.41 |
| with ERM | 512 | 1,000 | 83.70 |
|  |  | 2,000 | 89.36 |
|  |  | 4,000 | 92.86 |

Consequently, SSAlign-prefilter selects the 512-dimensional embeddings processed by ERM as the feature representation for proteins.

### SSAlign Database and WebServer

To address high memory demands in storing protein embedding representations, we implemented an efficient optimization solution using the FAISS library. FAISS is specifically designed for scalability and efficiency, capable of processing billion-scale vector databases with optimized CPUs/GPUs performance while significantly reducing time consumption. For the AFDB50 database containing 50 million sequences, FAISS successfully achieved substantial time reduction (refer to “Order-of-magnitude acceleration in structure search” subsections). For SAligner’s requirement to rapidly retrieve target protein 3Di sequences from large databases, we created an indexed 3Di sequence database for all proteins in AFDB50, which dramatically reduced both memory usage and execution time. We constructed complete SSAlign Database for Swiss-Prot, SCOPe40 and AFDB50, comprising two components:

- Vector Index Database (for SSAlign-prefilter): Built using the FAISS library.
- 3Di Sequence Database (for SAligner rapid retrieval): Enables efficient target protein 3Di sequence searches, Maintains low memory footprint.

Both components were designed with low memory consumption objectives. We have developed a web server for SSAlign, accessible at [Web server](http://bioinfo.isyslab.info/ssalign/search/)(Fig. 8). Users submit search requests via a browser by selecting the target database and retrieval size, and providing query structures either by pasting PDB text or uploading structure files. The Flask-based service validates inputs and creates a unique Job ID, then dispatches search jobs asynchronously through a Redis/Celery queue, where backend workers invoke the SSAlign engine for execution. Job status and results are organized by Job ID and presented on a dedicated results page; optional email notification provides a result link upon completion, enabling convenient asynchronous and batch usage for long-running searches. Additionally, through the web server, you can download the original data involved in this study as well as the [SSAlign Database](http://bioinfo.isyslab.info/ssalign/download/section/ssalign/).

**Figure 8.**
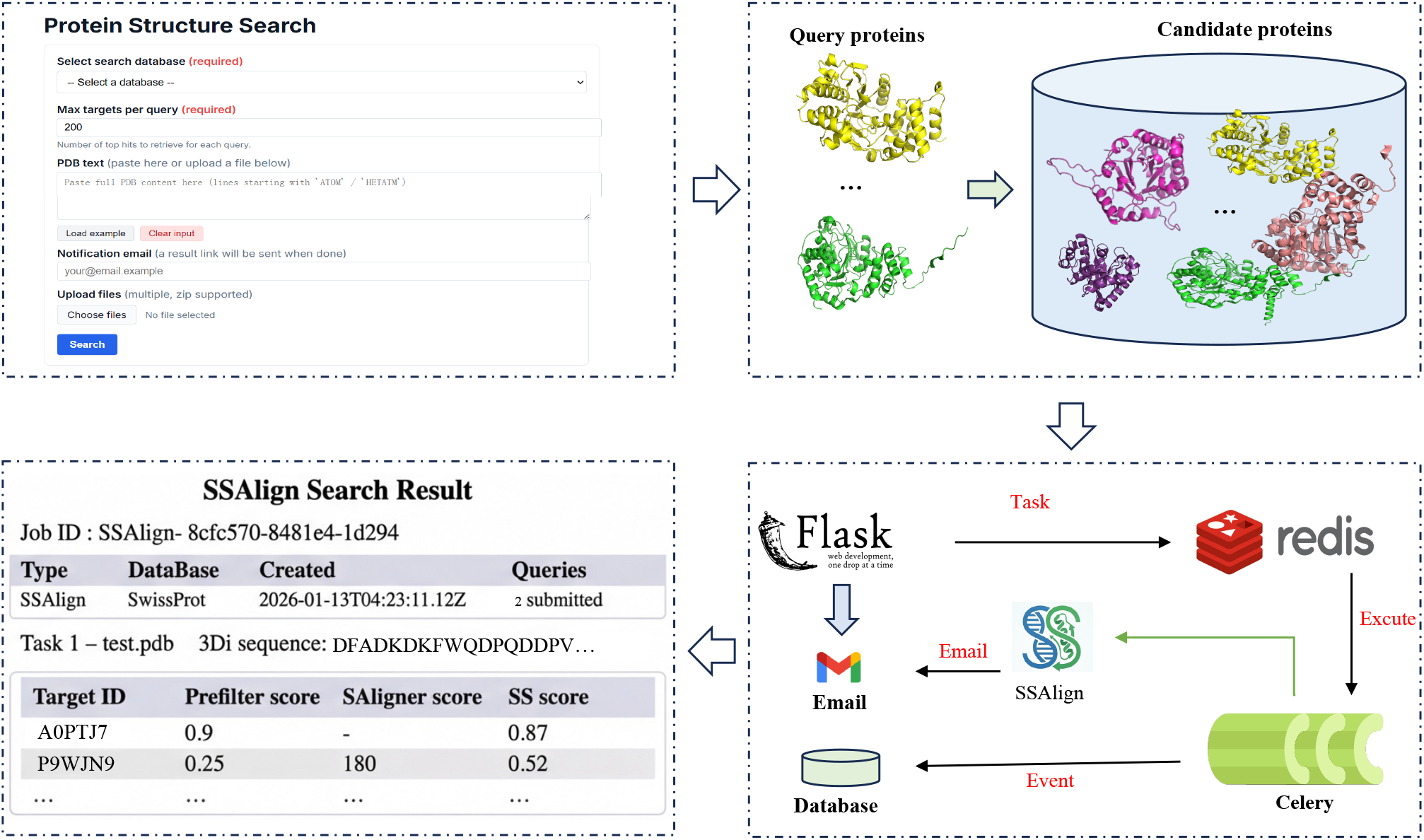
Workflow of the SSAlign web server. Users submit query structures via a web browser. The Flask-based web application validates the input, generates a unique job ID, and dispatches the search task asynchronously through Redis and Celery. Results are subsequently returned on a dedicated job page, with optional email notification.

### SS-score-predictor

The core of structural similarity computation lies in effectively comparing and evaluating the similarity between protein structures. The similarity metric in the SSAlign-prefilter stage relies on vectorized retrieval, typically using cosine similarity, but lacks biological significance for protein homology search. Traditionally, TM-score^23^ serves as an important standard for evaluating protein structural similarity. It is calculated based on distance variations of C_*α*_ atoms while considering protein size and shape, demonstrating good compatibility with protein structures of different lengths. However, TM-score computation generally requires comparing entire structures, which consumes substantial computational resources in large-scale alignments. The avg_TM-score is defined as the average of the normalized TM-scores for both query and target chains:

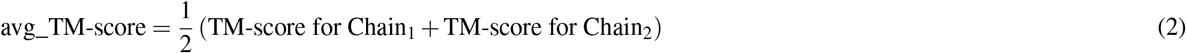

To enable rapid and effective assessment of structural similarity between proteins, SSAlign employs an evaluation metric called the SS-score.The SS-score-predictor estimates the avg_TM-score based on vector retrieval metrics (Cosine_Similarity) from the SSAlign-prefilter stage. Supplementary Table 12 presents the fitting parameters of the SS-score-predictor on both Swiss-Prot and SCOPe40 datasets. Using the Swiss-Prot dataset as an example, the linear regression yields:

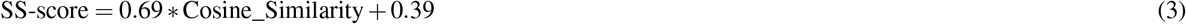

The Pearson correlation coefficient between the SS-score and avg_TM-score is 0.69. Unlike traditional tools like Foldseek that focus on local similarity, the SS-score shares the same design philosophy as TM-score by emphasizing global structural similarity. The SS-score demonstrates a strong positive correlation with TM-score(Supplementary Figures 1-2). This indicates that the SS-score effectively measures overall protein structural similarity rather than merely focusing on local matches between individual amino acid residues.

### Numba-accelerated SAligner

The Needleman-Wunsch algorithm represents a widely used global alignment method in sequence comparison^24,25^. As a dynamic programming approach, it identifies the optimal alignment between two sequences (e.g., proteins or DNA) by maximizing their similarity (or minimizing differences). Sequence alignment involves comparing multiple base/residue arrangements to reveal sequence similarity and establish homology. This methodology stems from the fundamental biological principle that sequence determines structure, and structure determines function - treating nucleic acids and protein primary sequences as character strings enables detection of functional, structural, and evolutionary information through similarity analysis. Compared to local alignment algorithms (e.g., Smith-Waterman^26^), global alignment better captures complete evolutionary relationships and structural contexts.

For protein sequence alignment, substitution matrices like BLOSUM62^27,28^ or PAM250^29^ conventionally quantify the likelihood/frequency of amino acid replacements. Leveraging Foldseek’s advanced structural encoder that discretizes protein structures into 3Di sequences, we implemented a Needleman-Wunsch variant for 3Di sequence alignment using the structural substitution matrix, which quantifies replacement probabilities between different 3Di alphabets. To optimize modern multi-core architectures, we pre-compiled the algorithm using the Numba compiler and enabled parallel execution of multiple alignments (default: 64 concurrent tasks). Supplementary Figures 1-2 illustrates the SAligner can effectively identify high avg_TM-score targets through the SAligner Score.

The Numba compiler employs Low-Level Virtual Machine (LLVM) and NVIDIA Virtual Machine(NVVM) technologies^30^. While LLVM translates interpreted languages like Python/Julia directly into CPU-executable machine code, NVVM extends this capability for GPU architectures. This compilation approach achieves a 100-fold speed improvement over interpreted execution. Taking the alignment of two 3Di sequences with length 1000 as an example, the computational time consumption is shown in Supplementary Table 13.

### Hyperparameter Configuration

The two-stage design of SSAlign allows customizable sensitivity control through parameter adjustment in each stage. Key parameters are described as follows:

- **db**: Specifies the target database to search. Currently supported: `Swiss-Prot,SCOPe40,AFDB50`.
- **querypdbs**: Comma-separated list of query structure files (e.g., `.cif`). Each query is searched independently against the selected database.
- **prefilter_target**: Number of candidates retained after the FAISS-based SSAlign-prefilter stage (default: 2000). Larger values improve recall but increase runtime and memory usage. In `mode=1`, this pool is used to construct the candidate list for the second-stage rescoring.
- **prefilter_threshold**: Score cutoff used to trigger second-stage SAligner rescoring (default: 0.3). Candidates with prefilter scores below this threshold are treated as marginal and are re-ranked by SAligner to improve reliability, as low prefilter scores tend to be less correlated with structural similarity.
- **max_target**: Maximum number of final homologs reported per query (default: 1000). The output is truncated to this limit after all ranking/rescoring steps.
- **mode**: Execution mode. `0` runs SSAlign-prefilter only; `1` runs the full two-stage pipeline (prefilter + SAligner rescoring).
- **prefilter_mode**: Backend for the FAISS prefilter stage (default: `cpu`). `cpu` performs retrieval on CPU, while `gpu` offloads retrieval to available GPUs and shards the FAISS index across multiple GPUs when applicable.
- **out_dir**: Output directory used to store search results and intermediate artifacts.
- **nproc**: Number of CPU threads/processes used for parallel execution in the SAligner stage (default: 64).
- **cuda_device**: CUDA device identifier for SaProt inference (e.g., `cuda:0`). This controls where the embedding/model forward pass is executed.

Supplementary Figure 3 illustrates the impact of the *prefilter_threshold* parameter on the precision–recall trade-off. This analysis validates the critical role of threshold selection in balancing detection sensitivity and alignment quality. Results obtained with lower thresholds require correction through SAligner’s realignment and rescoring to ensure reliability.

### Datasets and Benchmark

The assessment of structural alignment accuracy is based on evaluating the degree of spatial consistency achieved when aligning protein pairs within their respective structures. This self-contained evaluation approach is unbiased as it does not rely on external classification systems that may be constructed using specific sequence or structural alignment tools^31^. The TM-score serves as a global measure of alignment quality that is sensitive to coverage, ranging from 0 to 1, where 1 represents perfect structural match. As a key indicator of alignment accuracy, the TM-score range of [0.5, 1] is particularly important as it reflects the sensitivity of detecting structurally similar proteins^32^. The TM-score can be normalized by the length of either protein in the pair. To ensure fair comparison, we use the average of both normalizations (avg_TM-score) to mitigate biases from extreme length differences, providing a more robust assessment of structural similarity. The RMSD, normalized by the number of aligned residue pairs, provides a complementary measure of spatial proximity that effectively captures local alignment accuracy by quantifying the average distance deviation between corresponding atomic positions in the two protein structures.

For our Swiss-Prot benchmark (using protein structures from AlphaFold Database^33^), we randomly selected 100 query proteins and evaluated tool performance using both TM-score and RMSD metrics calculated by TM-align. The evaluation procedure consisted of:

1. Executing structural alignment tools.
2. Calculating TM-scores for all generated alignments using TM-align.
3. Ranking results by each tool’s native metrics (e.g., E-value, SS-score).
4. Additionally ranking alignments by TM-align calculated avg_TM-score.
5. Finally, computing cumulative avg_TM-score and RMSD values for both ranking methods.

This approach provides a comprehensive assessment of both alignment accuracy and retrieval efficiency. The SCOPe dataset^34,35^ provides a carefully curated hierarchical classification of protein structural domains based on structural and evolutionary relationships. We used the SCOPe40 dataset, containing 11,211 non-redundant protein sequences clustered at 40% sequence identity from SCOPe 2.01. Leveraging the known hierarchical relationships, we defined matches within the same family or superfamily^36^, or those with avg_TM-score ≥ 0.5 up to the 1st FP, as true positives (TPs), while matches not sharing the same fold with avg_TM-score *<* 0.5 were classified as false positives (FPs). Benchmarking was performed by ranking results according to each tool’s native metrics and measuring, for each query, the fraction of TPs retrieved before encountering the first FP. We conducted an all-versus-all search after excluding 13 problematic proteins to ensure clean evaluation.

For large-scale performance testing, we utilized the AFDB50 database containing 53,665,860 protein structures. This included comprehensive speed benchmarking and construction of complete FAISS index files to enable efficient database searches. Similarly, we conducted the same benchmarking procedure on AFDB50 as used for the Swiss-Prot dataset.

## Supporting information

Supplementary Materials for SSAlign

## Data and code availablity

All datasets, intermediate processing results, and prebuilt indices are publicly available at [SSAlign web server](http://bioinfo.isyslab.info/ssalign/download/section/ssalign/). The source code for SSAlign is available under an open-source license at https://github.com/ISYSLAB-HUST/SSAlign.

## Acknowledgements

This work was supported by National Natural Science Foundation of China under Grant 62172172, Hubei Provincial Natural Science Foundation of China under Grant 2025AFB159, and the Postdoctoral Fellowship Program of CPSF under Grant Number GZC20240545.

## Author contributions statement

L.W. and X.Z. contributed equally to this work and were responsible for the conceptualization, methodology, and initial drafting of the manuscript. Y.W. and Z.X. supervised the project, provided critical revisions, and contributed to data analysis. Z.X. coordinated the research activities, acquired funding, and finalized the manuscript. All authors reviewed and approved the final version of the manuscript.

