## Supplementary Materials for SSAlign for "SSAlign: Ultrafast and Sensitive Protein Structure Search at Scale"

### Simple fold protein cases

We identified 13 simple fold protein protein cases (though not peptides) in the SCOPe40 dataset that Foldseek failed to retrieve. As demonstrated in Table 1, SSAlign exhibits robust performance at both family and superfamily levels within the SCOPe40 dataset. These results establish SSAlign's superior performance in identifying challenging structural relationships that elude existing detection methods. Tables 2 presents the retrieval performance of SSAlign for the seven antimicrobial peptide (AMP) cases discussed in the main text, showing that Foldseek failed to match even themselves (denoted by "/" in the table), where "1" indicates only self-matching. In contrast, SSAlign successfully detected most homologous pairs of these AMPs, with the average avg_TM-score of the top 10 results listed. Tables 3–9 present seven representative AMPs identified in Swiss-Prot. For each peptide, we show the top 10 retrieval results obtained using SSAlign, with structural similarity evaluated by TM-align based on RMSD and average TM-score calculations. Table 10 summarizes additional AMP cases from Swiss-Prot, where the 3Di sequences encoded by Foldseek exhibit distinctive characteristic patterns.

**Supplementary Table 1.** Identification of simple fold protein in SCOPe40

| | d1dpjb_ | d2e74d | d1q90g | d1xoua_ | d1ehkc_ | d1l2paa_ | d2ciob_ | d1q90m_ | d1ehkb2 | d1jb0x_ | d1jb0m_ | d1q90a3 | d1rzhh2 |
|---|---|---|---|---|---|---|---|---|---|---|---|---|---|
| Super Family | 1 | 4 | 1 | 2 | 1 | 1 | 3 | 2 | 4 | 1 | 1 | 1 | 3 |
| Family | 1 | 4 | 1 | 1 | 1 | 1 | 1 | 2 | 4 | 1 | 1 | 1 | 3 |
| TM-align | / | / | / | / | / | / | / | / | / | / | / | / | / |
| SSAlign-prefilter-500 | 1 | 4 | 1 | 2 | 1 | 1 | 1 | 2 | 2 | 1 | 1 | 1 | 3 |
| SSAlign | 1 | 4 | 1 | 2 | 1 | 1 | 1 | 2 | 2 | 1 | 1 | 1 | 3 |

**Supplementary Table 2.** Antimicrobial peptide cases in Swiss-Prot

| ID | 3Di sequences | Foldseek | SSAlign |
|---|---|---|---|
| P84381 | DPVPVVVVVVCVVVCVVPVVPPDDPPD | / | 0.66 |
| P84382 | DVVVVVVVVVVVCVVVVVCCVVDVD | 1 | 0.80 |
| P84383 | DVCVVVVVVVVCCVVPVVVVVVPPD | 1 | 0.58 |
| P84384 | DVVVVVVVVCVVPVVVCVVPVD | 1 | 0.53 |
| P84385 | DCPVVVVVVVVVVVCVVVVVVD | / | 0.72 |
| P84386 | DVVVVCVVVVVVVVVVVD | / | 0.69 |
| P84387 | DVCPVVVVVVVVVVVVVD | 1 | 0.50 |

**Supplementary Table 3.** P84381

| target | RMSD | avg_TM-score |
|---|---|---|
| P84381 | 0.0 | 1.0 |
| P84386 | 0.57 | 0.614 |
| C0HKN4 | 0.66 | 0.689 |
| C0HKN3 | 0.59 | 0.663 |
| C0HKL4 | 0.53 | 0.709 |
| C0HKP4 | 0.76 | 0.573 |
| P82393 | 0.55 | 0.655 |
| C0HKL2 | 0.64 | 0.544 |
| C0HKN5 | 0.64 | 0.552 |
| P85982 | 0.63 | 0.609 |

**Supplementary Table 4.** P84382

| target | RMSD | avg_TM-score |
|---|---|---|
| P84382 | 0.0 | 1.0 |
| C0HKM0 | 0.54 | 0.785 |
| C0HKP5 | 0.65 | 0.764 |
| C0HKM1 | 0.64 | 0.785 |
| C0HKM2 | 0.47 | 0.814 |
| C0HKP7 | 0.46 | 0.821 |
| C0HKP6 | 0.55 | 0.785 |
| P84384 | 1.49 | 0.476 |
| P86416 | 0.77 | 0.597 |
| P84925 | 1.08 | 0.594 |

**Supplementary Table 5.** P84383

| target | RMSD | avg_TM-score |
|---|---|---|
| P84383 | 0.0 | 1.0 |
| C0HKM8 | 1.34 | 0.576 |
| C0HLC7 | 1.62 | 0.505 |
| C0HLC5 | 1.53 | 0.510 |
| P84923 | 1.61 | 0.511 |
| C0HLC8 | 1.62 | 0.524 |
| C0HKM9 | 1.37 | 0.563 |
| C0HKP3 | 0.65 | 0.553 |
| C0HK89 | 1.52 | 0.575 |
| C0HKN2 | 1.41 | 0.557 |

**Supplementary Table 6.** P84384

| target | RMSD | avg_TM-score |
|---|---|---|
| P84384 | 0.0 | 1.0 |
| P84383 | 0.74 | 0.580 |
| C0HKP5 | 1.56 | 0.479 |
| C0HKM1 | 1.52 | 0.491 |
| C0HKM2 | 1.54 | 0.491 |
| C0HKP7 | 1.56 | 0.488 |
| P84382 | 1.49 | 0.475 |
| C0HK85 | 1.5 | 0.492 |
| C0HKM0 | 1.68 | 0.479 |
| C0HKN2 | 1.37 | 0.485 |

**Supplementary Table 7.** P84385

| target | RMSD | avg_TM-score |
|---|---|---|
| P84385 | 0.0 | 1.0 |
| C0HKL7 | 0.46 | 0.727 |
| C0HKN9 | 0.44 | 0.736 |
| C0HKP2 | 0.33 | 0.755 |
| C0HK88 | 0.37 | 0.758 |
| C0HKP1 | 0.42 | 0.700 |
| C0HKP0 | 0.69 | 0.599 |
| C0HK87 | 0.61 | 0.630 |
| C0HKP4 | 0.4 | 0.677 |
| C0HKP3 | 0.46 | 0.663 |

**Supplementary Table 8.** P84386

| target | RMSD | avg_TM-score |
|---|---|---|
| P84386 | 0.0 | 1.0 |
| C0HKN4 | 0.38 | 0.692 |
| C0HKN3 | 0.23 | 0.831 |
| P84381 | 0.57 | 0.613 |
| C0HKL4 | 0.44 | 0.615 |
| C0HK91 | 0.43 | 0.670 |
| P0DTV6 | 0.32 | 0.679 |
| C0HKP4 | 0.5 | 0.575 |
| P86281 | 0.47 | 0.664 |
| P85443 | 0.6 | 0.564 |

**Supplementary Table 9.** P84387

| target | RMSD | avg_TM-score |
|---|---|---|
| P84387 | 0.0 | 1.0 |
| C0HKM8 | 2.46 | 0.442 |
| C0HKM9 | 1.98 | 0.460 |
| C0HKM7 | 1.75 | 0.407 |
| B3EWT9 | 1.83 | 0.450 |
| C0HK89 | 2.12 | 0.469 |
| B3EWT6 | 1.98 | 0.405 |
| P84381 | 0.68 | 0.549 |
| C0HKQ6 | 0.99 | 0.365 |
| C0HKN1 | 1.09 | 0.440 |

**Supplementary Table 10.** Antimicrobial Peptide's 3Di Sequences in Swiss-Prot

| ID | 3Di Sequence |
|---|---|
| B3A0L2 | DCPDPPQDPVPDDPPDDDD |
| B3A0L3 | DPPDDPDDPPDPDDDPPDDD |
| P0CI89 | DVVVVVVVVVVVVVVVVVVVVVVVVPPCVVVVVVVVVVVVVVVVVVCVVPVDPVVCPVVVVVVVVVVVVPDDDD |
| P0CI90 | DVVVVVVVVVVVVVVVVVVVVVVVVVVVVVVVVVVVVVVVVVVVVVVCVVVVVVVVVVVVVVVVVVVPDDDD |
| G8YYA6 | DPVVVVVVVVVVVVVVCVVCPPPCVPPVVVVVVVVVVVVVVVVVVVVVVVVVVVVVVVVVVVVVVVVVVPDDPD |
| G8YYA5 | DVVVVVVVVVVVVVVVVVVVVCCVVPVVVVVVVVVVVVVVVVVVVVVVVVVVVVVVVVVVVVVVVVVVVVVPDDPD |
| C0HKQ5 | DVVVVVVVVVVVVVVVD |
| C0HKQ6 | DCVVVVVVVVVVVVPPD |
| A0A0A1I6N9 | DPVVVVVVVVVVVVVVVVVVVCCCVPPVVVVVVVVVVVVVVVVVVVVVVVVVVVVVVVVVVVVVVVVVVVPDDDD |
| A0A0A1I6E7 | DVVVVVVVVVVVVVVVVVVVVVCCVPPPVVVVVVVCVVVVVVVVVVVVVVVVVVVVVVVVVVVVVVVVVPDDPD |
| L0GAZ8 | DVVVLVVVLVVVVVVVVQVVVVVSVVVVVVVVCVVCDPVVVVVVVVVVVVPVPDCDVVNVVVNVVVVD |
| L0GCI6 | DVVVVVVVVVVVVVVVVQVVDPVSVVVVVVVVCVVCDPVVVVCVVVVVVCVVPPPDVVVVVVVVVVVD |
| L0GBQ6 | DPVVVVVVVVVVVVVPPDPPDVVVVVVVVVVVVVCVCPDPVVVVVVVVLQVLLVQLCVVVVHHSVVLVVPDPVVSVVSVVD |
| L0GCV8 | DVVVVVVVVVVVVVVVCCVVCVPPVVVVVVVVVCVVCVVVVVVVVVVVVVVVVVVVVVVVVVVVVVVVVVVVVVVVVVVVVVD |
| P0DMI9 | DDPVVVVVVVVVVVVVVVVVVVVVVVVVVVVVVCVVVPPCVVVVVVVVVVVVVVVVVVCVVDPPPPDDPPPDDD |
| P0DMI7 | DDPVVVVVVVVVVVVVVVVVVVVVVVVVVVVVVVVVVPDPDVVVVVVVVVVVVVVVVVVVPDPPPDDDDDDDDD |
| P0DMJ0 | DDPVVVVVVVVVVVVVVVVVVVVVVVVVVVVVVVVVDDDDDVVVVVVVVVVVVVVVVVVVPPDPPPDDDPPDDD |
| P0DMI8 | DCPVVVVVVVVVVVVVVVVVVVVVVVVVVVVVVCVVPVPDDPVVVVVVVVVVVVVVVVVVPDPDPDPDPDDDDD |
| P0CH48 | DDDDPPPPDDDDDDDDDPDPD |
| P0CH49 | DVVVVVVVVVVVVVVVVVVVVVPPD |
| S6CWV8 | DPVVVVVVVVVVVVVVVVVVVVVVVVVCVVVVVVVVVVVVVVVVVVVVVVVVVVVVVVVVVVVVVVVVVPDPPD |
| S6D3A7 | DPVVVVVVVVVVVVVVVVVVVVVVVVVVVVVVVVVVVVVVVVVVVVVVVVVVVVVVVVVVVVVVVVVVVPDDPD |
| G1FE62 | DDPVVVVVVVVVVVVVVVVVVCCVVPPVVVVVVVVVVVVVVVVVVVVVVVVVVVVVVVVVVVVVVVVVVVVVPDDDD |
| P84385 | DCPVVVVVVVVVVVCVVVVVVD |
| P84384 | DVVVVVVVVCVVPVVVCVVPVD |
| P84382 | DVVVVVVVVVVVCVVVVVCCVVDVD |
| P84381 | DVCPVVVVVVVVVVVVVD |
| P84387 | DPVPVVVVVVCVVVCVVPVVPPDDPPD |
| P84383 | DVCVVVVVVVVCCVVPVVVVVVPPD |
| P84386 | DVVVVCVVVVVVVVVVVD |
| P39080 | DVLVVQLVVVVVVPVVSNVVVVVD |

### Structure-aware Protein language model

We chose SaProt due to its excellent performance in protein representation and its ability to capture deep patterns and features. For the embedding layer selection, we randomly sampled a small subset from the Swiss-Prot dataset for validation. In this validation, we compared the 33rd layer embeddings with the average embeddings of the 1st and 33rd layers(Table 11), calculating the overlap rate between the top-k results screened by the SSAlign-prefilter stage and those of Foldseek. The results showed that the 33rd layer embeddings exhibited superior representational capability. In this study, we selected SaProt_650M_AF2 as it demonstrated marginally superior performance in our evaluations.

**Supplementary Table 11.** Overlap rates with Foldseek at different layer

| layer | Top-k | Overlap Rate (%) |
|---|---|---|
| 33 | 1,000 | 85.36 |
| | 2,000 | 89.94 |
| | 4,000 | 92.41 |
| (33+1)/2 | 1,000 | 83.87 |
| | 2,000 | 88.89 |
| | 4,000 | 91.59 |

### SS-score-predictor and SAligner

Table 12 presents the fitting parameters of the SS-score-predictor when SSAglin employs on both Swiss-Prot and SCOPe40 datasets. Figs. 1–2 illustrate two representative cases that demonstrate the reference value of SS-score predictions generated by the SS-score-predictor and validate the effectiveness of SAligner(For the low-threshold results filtered by the SSAlign-prefilter stage, the second-stage SAligner can effectively identify high avg_TM-score targets through the SAligner Score, thereby compensating for the limitations of SSAlign-prefilter).

Table 13 demonstrates the accelerated computational performance of our Numba-accelerated SAligner implementation.

**Supplementary Table 12.** SS-score-predictor parameters

| Dataset | $a_1^{[1]}$ | $a_2^{[1]}$ |
|---|---|---|
| Swiss-Prot | 0.69 | 0.39 |
| SCOPe40 | 0.59 | 0.42 |

**Supplementary Table 13.** Time Consumption for Aligning Two 3Di Sequences of Length 1000

| Non-accelerated SAligner(Seconds) | Biopython(Seconds) | SAligner(Seconds) |
|---|---|---|
| 0.48620128631591797 | 0.006602048873901367 | 0.005364656448364258 |

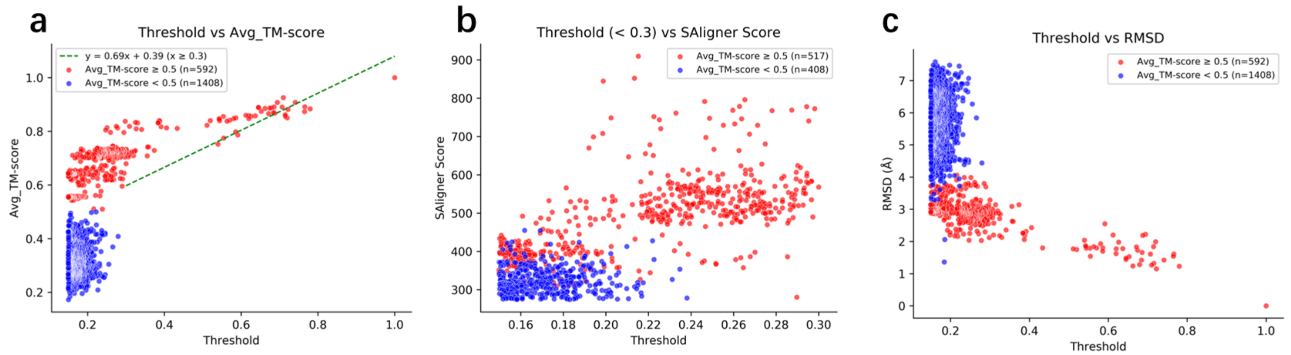


**Supplementary Figure 1.** The SSAlign results for query F1Q575.Subfigures (a) and (c) fully validate that the scores from the SSAlign-prefilter stage exhibit a high correlation with protein structural similarity metrics. An appropriate threshold ensures the high accuracy of SSAlign-prefilter, while the curve in subfigure (a) demonstrates that the SS-score predicted by the SS-score predictor is strongly correlated with the avg_TM-score, confirming that the SS-score effectively measures protein structural similarity. Figure (b) illustrates how the re-ranking function of SAligner correctly filters potential hits with avg_TM-score $\geq$ 0.5 from low-threshold candidates.

---

### prefilter_threshold

Fig. 3 illustrates the effect of different `--prefilter_threshold` values on retrieval performance under a fixed `--prefilter_targe`
`2000` on the target dataset.

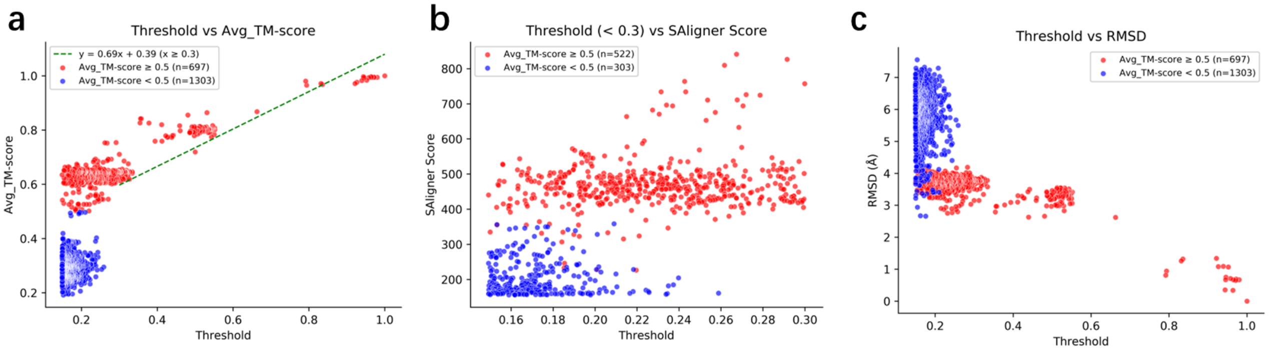


**Supplementary Figure 2.** The SSAlign-prefilter results for query Q99VP7.

---

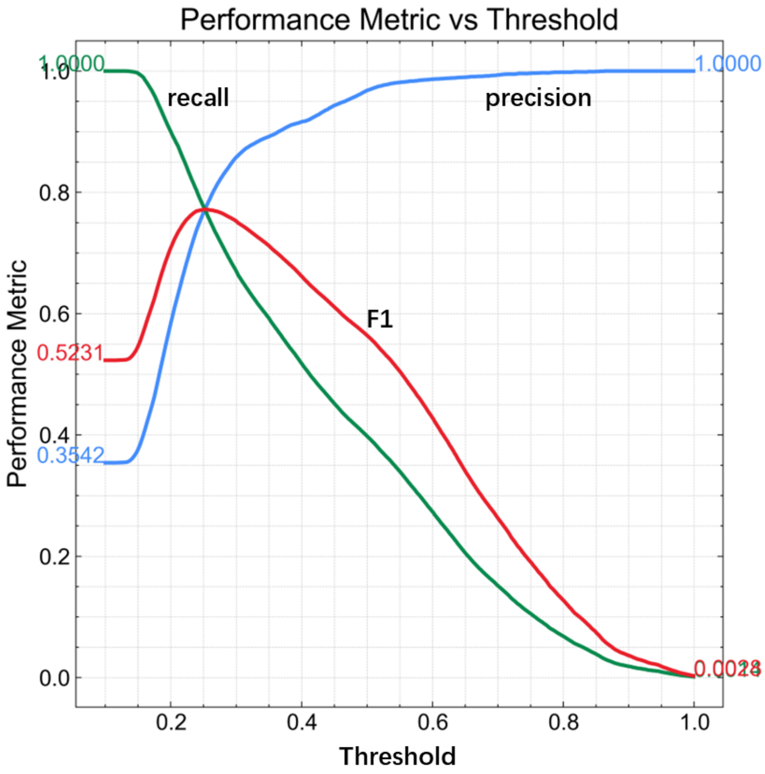


**Supplementary Figure 3.** **Optimal `−prefilter_threshold` values across embedding dimensions**. For the default dimensional embeddings (`−prefilter-target 2000`), we recommend `−prefilter_threshold` = **0.3**
